# J-domain proteins bridge the PKL and ATRX complex to transcription factors to fine-tune gene expression via H3.3 deposition

**DOI:** 10.64898/2026.08.25.747133

**Authors:** Shuya Wang, Colette L. Picard, Zhongshou Wu, Yan He, Alexander Barinsky, Evan K. Lin, Russell Chuang, Lu Li, Jihui Sha, James Wohlschlegel, Suhua Feng, Steven E. Jacobsen

## Abstract

Epigenetic modifications, including histone modifications and DNA methylation, direct gene expression programs during the growth and development of eukaryotic organisms. In *Arabidopsis thaliana*, the chromatin remodeler PICKLE (PKL), a homolog of animal CHD3, plays a critical role during these processes. Previous studies of PKL have painted a complex picture of its function, including both activating and repressive roles, however, how PKL can control both transcriptional silencing and activation is unknown. We have identified a group of J-domain-containing proteins (DNAJs), usually known for their roles in protein folding, that guide PKL recruitment to various gene promoters by bridging PKL to transcription factors. Mutation of PKL’s DNAJ-interacting domain disrupts PKL association with chromatin. Once recruited to transcription factor-bound sites, PKL coordinates with the histone chaperone ATRX to deposit HISTONE3.3 (H3.3) at targeted loci, which also negatively affects the accumulation of H3 lysine 27 trimethylation (H3K27me3) at promoters. We found that transcriptional outcome of PKL binding depends not only on the changes in H3.3 occupancy but also on the pre-existing chromatin context at PKL-targeted sites. Our findings outline a new mechanism for CHD3 chromatin remodeler recruitment and function.

## Introduction

The epigenome is dynamically modified to alter gene expression as needed for different developmental stages and cell types. During this process, nucleosome positioning and histone and DNA modifications coordinate with a host of epigenetic regulators and transcription factors to achieve precise control of transcription [1–4]. Chromatin remodelers are ATP-dependent proteins that modulate nucleosome positioning, influence histone modifications, and control transcription during development [4]. The CHD (chromodomain helicase DNA-binding) family of chromatin remodelers, conserved in many eukaryotes including plants and animals, is one example [5]. In mammals, CHD3, a component of the NuRD complex, is recruited in part by the reader proteins MBD2 or MBD3 to alter nucleosome structure to fine-tune gene expression [6, 7]. For example, the NuRD complex restrains inappropriate lineage development and is essential in neural stem/progenitor fate decisions during brain development [8, 9]. PKL, a well-studied CHD3 protein in *Arabidopsis*, is similarly important for proper development. PKL has been identified as a regulator of embryonic identity during the seed-to-seedling transition and the juvenile-to-adult transition, as well as a regulator of root meristem identity [10–14]. Transcription factors VAL1 and VAL2 have been reported to guide PKL recruitment to seed maturation genes [15, 16]. Transcription factors such as TCPs, PIFs, and HY5 also appear to facilitate PKL recruitment to some of its euchromatic targets but can only account for a subset of PKL binding sites [16, 17]. It is unclear how these different recruitment routes vary across tissues and developmental stages. Furthermore, accumulating evidence suggests that PKL contributes to both transcriptional silencing and activation [11, 16, 18]. While PKL has been reported to promote the spreading of the repressive modification H3K27me3 at some pre-existing Polycomb domains, it has also been reported to support deposition of the active modification H3K4me2 and maintain permissive chromatin at some euchromatic genes [16, 19]. These apparently opposing functions have led to the view that PKL acts in a highly context-dependent manner. Yet the mechanistic basis of this remains poorly understood [15, 16, 19].

J-domain-containing proteins (DNAJs), broadly required for protein homeostasis, plant growth and development, are also involved in both transcriptional silencing and activation [20, 21]. DNAJs, also known as Hsp40s, are HSP70-associated co-chaperones that regulate protein folding, complex assembly, and proteostasis [22–26]. Recent studies have expanded this canonical view by implicating J-domain proteins in transcriptional activation and repression [27, 28]. For example, the DNA methylation reader complex containing SUVH1/SUVH3 utilizes DNAJ1 and DNAJ2 to enhance expression of genes nearby methylated transposons, indicating that J-domain proteins can interact with epigenetic ‘readers’ to carry out a specific regulatory function at a specific subset of genomic loci [27, 29, 30]. Similarly, the MBD5/MBD6 complex silences DNA-methylated genes and transposable elements [31] via the J-domain protein SILENZIO [28]. These findings suggest that J-domain proteins can act as chromatin-associated cofactors that help regulate local epigenetic states.

Here, we identify a novel group of DNAJs that guide PKL chromatin localization and function by anchoring PKL to various transcription factors, thereby directing PKL to specific genes. Loss of the DNAJ-interacting domain of PKL severely disrupts PKL’s association with chromatin and results in mutant phenotypes mimicking the *pkl* mutant. Once recruited to target loci, which are mostly promoters and transcriptional start sites (TSSs), PKL interacts with the histone chaperone ATRX that deposits H3.3, thereby altering local nucleosome occupancy and composition. The eventual transcriptional outcomes following PKL and ATRX recruitment are determined by the combination of the preexisting local nucleosome landscape and changes in H3.1/H3.3 ratio. Our results provide a mechanistic explanation for PKL’s highly context-dependent role in transcriptional regulation via a novel DNAJ-mediated interaction, extend our understanding of PKL-mediated transcriptional regulation, and highlight a new paradigm in CHD3 chromatin remodeler recruitment by DNAJ proteins.

## Results

### DNAJs are components of PKL and ATRX complexes

Transcription factors such as VAL1, VAL2, TCP4, PIFs, and HY5 have been shown to recruit PKL, but account for only a subset of PKL binding (Fig S1A-E) and were not consistently found associated with PKL across different studies [15, 16]. In addition, it’s difficult to determine whether these interactions are direct vs. indirect, since chromatin remodelers often associate with histone modifiers to influence transcription initiation and elongation [8, 32, 33]. To identify partners that guide PKL localization, we performed Immunoprecipitation Mass Spectrometry (IP-MS) using FLAG-tagged PKL driven by its endogenous promoter. PKL IP-MS identified four novel DNAJs as the top interactors (Fig 1A-C, Fig S1F-G), DNAJ40 (AT5G53150), DNAJ41 (AT2G25560), DNAJ68 (AT2G05230), and DNAJ84 (AT2G05250) [24]. DNAJ68 (AT2G05230) and DNAJ84 (AT2G05250) are tandemly duplicated and have identical amino acid sequences, so we refer to them collectively as DNAJ68 for simplicity. Peptides from the H3.3 chaperone ATRX were also highly enriched (Fig 1A and 1C, Fig S1F-G). ATRX is an SNF2-family chromatin remodeler implicated in replication-independent H3.3 deposition that regulates H3.1/H3.3 balance at expressed genes, including a subset of Polycomb-associated developmental genes [34–36].

**Figure 1.**
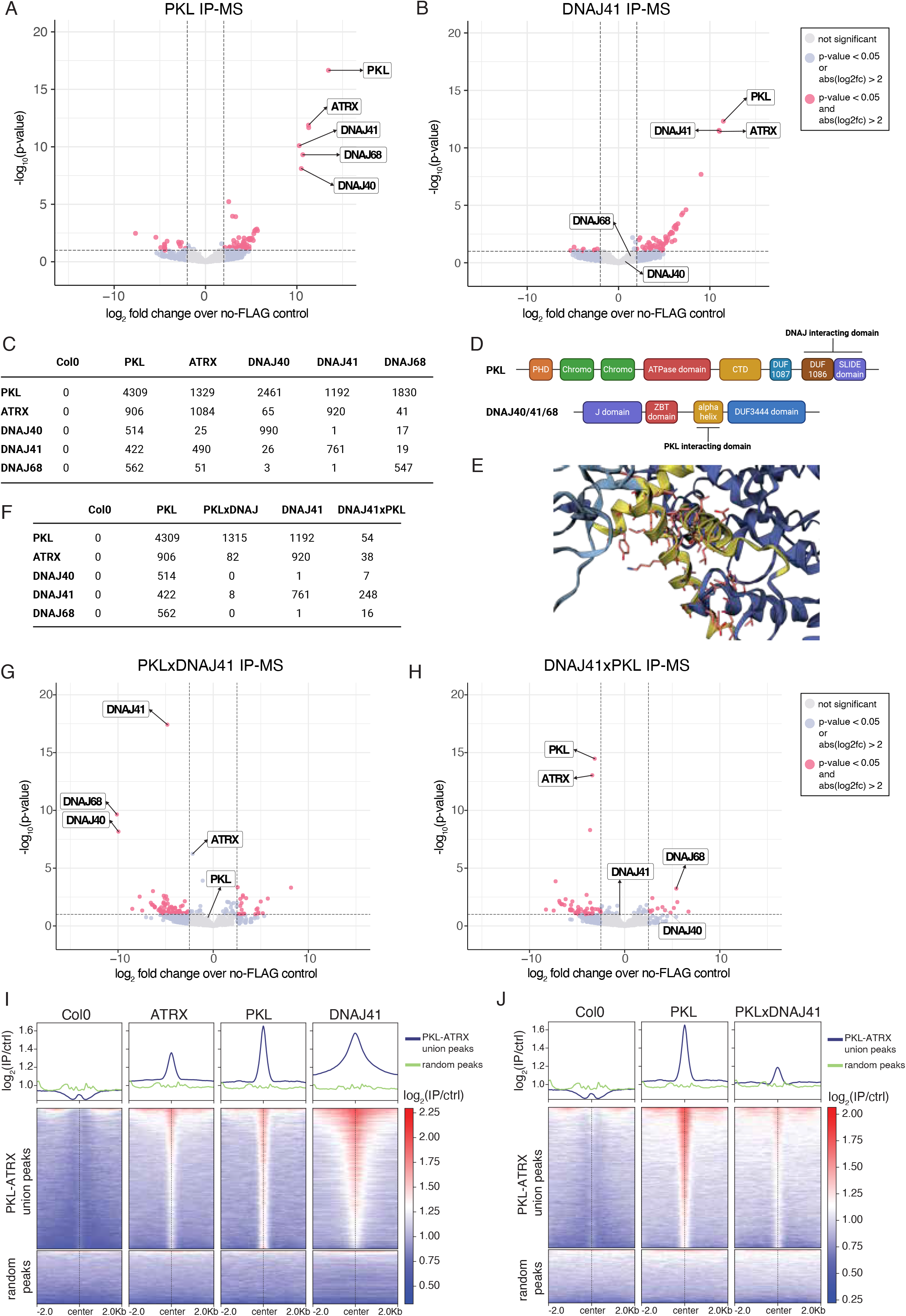
DNAJs are key components of the PKL-DNAJ-ATRX complex. Volcano plots showing protein enrichment from **A**. PKL IP-MS and **B**. DNAJ41 IP-MS. Pink dots are proteins significantly enriched or depleted compared to Col-0 control. **C**. Normalized protein enrichment summary of Col-0, PKL, ATRX, DNAJ40, DNAJ41, and DNAJ68 IP-MS. **D**. Key protein domains of PKL and DNAJ40/41/68. **E**. AF3 modeling of interaction between DNAJ41 and PKL. Light blue = DNAJ41, slate = PKL. Yellow indicates the interface secondary-structure elements highlighted on both proteins. Sticks are high-confidence interface residues. Pink = carbon atoms. Blue = nitrogen atoms. Red = oxygen atoms. The distances in Ångströms of the two salt bridges anchoring the core are: K358–E1050 (2.9 & 3.3 Å) and K360–E1064 (3.3 Å). **F**. Normalized enrichment summary of Col-0, PKL, PKLxDNAJ, DNAJ41, and DNAJ41xPKL IP-MS. Volcano plots showing normalized protein enrichment from **G**. PKLxDNAJ IP-MS normalized to PKL IP-MS and **H**. DNAJ41xPKL IP-MS normalized to DNAJ41 IP-MS. Metaplots and heatmaps of input-normalized ChIP-seq enrichment of **I**. Col-0, ATRX, PKL, and DNAJ41, and **J**. Col-0, PKL, and PKLxDNAJ over the union of PKL and ATRX ChIP-seq peaks vs. random peaks.

This suggests PKL may be involved in H3.3-related chromatin remodeling. To further validate these interactions, we FLAG-tagged the three DNAJs and ATRX and performed IP-MS. DNAJs, ATRX, and PKL consistently interacted with one another, suggesting that they form a stable complex (Fig 1B-C, Fig S1F-G, Fig S2A), which we named the PKL-DNAJ-ATRX complex. While PKL pulled down all three DNAJs equally, ATRX pulled down much more DNAJ41 than DNAJ40 and DNAJ68, suggesting preferential interaction between ATRX and DNAJ41. Although PKL was strongly associated with DNAJs and ATRX, we failed to detect significant enrichment for previously reported PKL interactors, including transcription factors VAL1/2, Histone Deacetylase 6 (HDA6), Polycomb Repressive Complex 2 (PRC2), or Trithorax1 (ATX1) [37–39]. Given that we performed IP-MS using floral tissues and previous reports of PKL IP-MS data were from seedlings, leaves, or suspension-cultured cells, it’s possible that these interactions are tissue-specific.

To understand whether and how PKL or ATRX interact with the DNAJs, we used AlphaFold 3 (AF3) to predict potential interaction interfaces. All DNAJs were predicted to use the same alpha helix, positioned between the J domain and the DUF344 domain, to interact with the SANT-SLIDE domain of PKL (Figure 1D-E, Fig S2B-D, Table S1-2), suggesting that each PKL can only bind a single DNAJ at a time. We created transgenes expressing FLAG-tagged PKL without the SANT-SLIDE domain, as well as FLAG-tagged constructs of each DNAJ without the alpha helix, and introduced these mutant versions into plants (Fig. 1D). We also tested a deletion of the alpha helix of DNAJ41, since DNAJ41 was the most enriched DNAJ from PKL and ATRX IP-MS. To confirm that these mutants did not destabilize the protein or cause protein degradation, we performed western blots of the truncated constructs of PKL and DNAJ41 alongside the full-length proteins. We observed no degradation for any of the mutants (Fig S2E). Removal of the SANT-SLIDE domain from PKL (hereafter named PKLxDNAJ) greatly reduced its interactions with all three DNAJs, while ATRX interaction was retained (Fig 1F and 1G, Fig S1G). Truncating the alpha helix of DNAJ41 (hereafter named DNAJ41xPKL) greatly reduced, but did not abolish, its interaction with PKL and ATRX (Fig 1F and 1H, Fig S1G), suggesting that additional protein domains in DNAJ41 may help mediate its interaction with PKL. Together, these data suggest that PKL forms a stable complex with ATRX and one of three DNAJs.

Next, we performed Chromatin Immunoprecipitation Sequencing (ChIP-seq) of 3xFLAG-tagged full-length DNAJ41, PKL, and ATRX driven under their endogenous promoter and found that DNAJ41, PKL, and ATRX colocalized at a subset of genomic regions, which were mainly promoters, transcription start sites (TSSs), and intergenic regions (Fig 1I-J, Fig S3A-B). These regions, defined as the union of PKL and ATRX ChIP-seq peaks, were marked by low nucleosome occupancy and high ATAC-seq signal, consistent with these sites being primarily accessible promoters (Fig S3C-D). We also performed ChIP-seq of PKLxDNAJ and DNAJ41xPKL, with full-length versions of PKL and DNAJ41 as controls.

PKLxDNAJ showed much weaker enrichment at PKL-targeted regions (Fig 1J), suggesting that abolishing the ability of the PKL to interact with the DNAJs impedes its ability to bind its target sites. On the other hand, DNAJ41xPKL had only moderately weaker binding to its target sites compared to wild-type DNAJ41 (Fig S3E), suggesting PKL recruitment is downstream of DNAJ’s binding to chromatin. Our results suggest that DNAJs help recruit the PKL-DNAJ-ATRX complex to its chromatin targets.

### DNAJ interactions are required for PKL-mediated gene regulation

Since PKLxDNAJ lost binding at PKL-complex binding sites, we examined whether the PKLxDNAJ construct can complement the *pkl* morphological and transcriptional phenotypes. The *pkl* mutant develops dark green rosette leaves with shortened petioles and delayed flowering, with more branching due to reduced apical dominance [10, 13, 40]. As a positive control, we first expressed full-length PKL driven under its endogenous promoter into the *pkl* mutant. Reintroducing full-length PKL into the *pkl* mutant rescued all the above phenotypes (Figure 2A). PKLxDNAJ, however, failed to complement PKL loss (Fig. 2A). We next tested whether PKLxDNAJ could rescue gene expression changes in the *pkl* mutant. RNA sequencing (RNA-seq) of floral tissues showed that gene expression changes in the *pkl* mutant were rescued by full-length PKL (Fig 2B-E). However, PKLxDNAJ was unable to rescue the transcriptional changes caused by PKL loss (Fig 2B-E).

**Figure 2.**
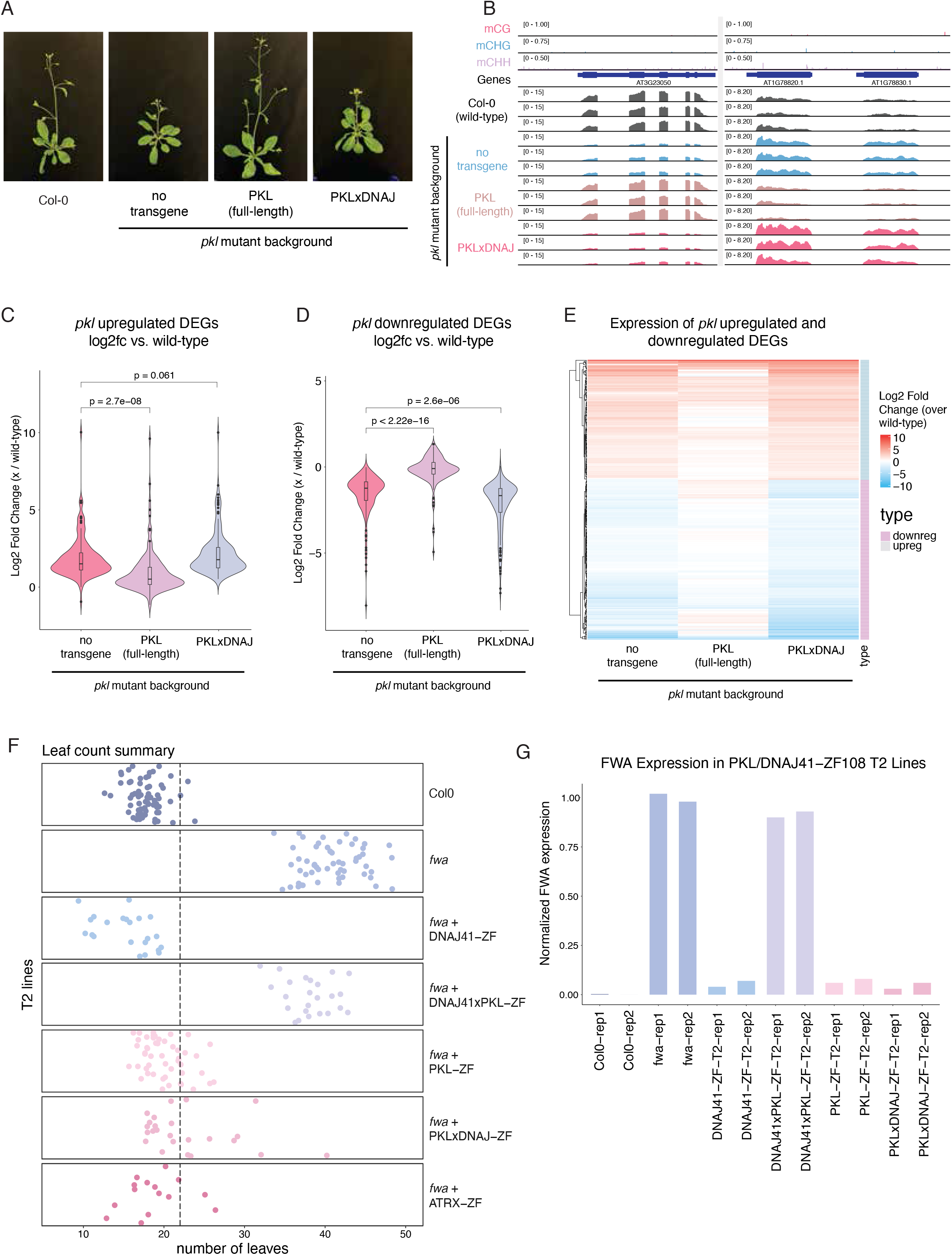
DNAJs are indispensable for PKL-mediated gene regulation. A. Images of 6-week-old wild-type (Col-0), *pkl*, PKLxDNAJ-Flag in *pkl*, and PKL-Flag in *pkl*. **B**. Genome browser images of RNA-seq tracks alongside DNA methylation and gene annotations at two PKL-regulated sites. Three replicates of RNA-seq shown for each genotype. **C-D**. Violin plots showing the log2 fold change (over Col-0 wild type) of gene expression at **C**. upregulated genes in *pkl*, or **D**. downregulated genes in *pkl* across *pkl*, PKL-Flag in *pkl*, and PKLxDNAJ-Flag in *pkl*. **E**. Heatmap showing the log2 fold change of gene expression at differentially expressed genes in *pkl*. **F**. Leaf count at bolting of Col-0, *fwa*, and T2 lines of DNAJ41-ZF108, DNAJ41xPKL-ZF108, PKL-ZF108, PKLxDNAJ-ZF108, and ATRX-ZF108. Cutoff lines indicate plants with ≤22 leaves. **G**. Bar plot summarizing RT-qPCR results from the Col-0, *fwa*, and corresponding T2 lines.

It is possible that deleting the DNAJ-interacting domain simply inhibits proper PKL protein folding and/or function. Since our data suggest that the DNAJs control recruitment of the PKL-DNAJ-ATRX complex, we tested whether tethering PKLxDNAJ to a specific locus could still cause silencing. We fused full-length PKL, PKLxDNAJ, and DNAJ41 with a zinc finger domain (ZF108) that binds the promoter of *FWA* [41, 42]. *FWA* encodes a transcription factor that is silent in wild-type due to DNA methylation present at its promoter [43]. In the epiallele mutant *fwa*, this DNA methylation is absent, and *FWA* becomes overexpressed, causing a late flowering phenotype that can be measured by counting the number of rosette leaves present after bolting [41, 42]. We transformed both PKL-ZF108 and DNAJ41-ZF108 into *fwa* and tested whether they were able to rescue the late flowering phenotype by counting the number of leaves after bolting. Per construct, we analyzed multiple T2 lines with verified protein expression. Both constructs were able to silence *FWA* transcription in *fwa* and restore normal flowering, as demonstrated by a decrease in leaf number (Fig 2F-G). PKLxDNAJ-ZF108 was also capable of silencing *FWA* and restoring early flowering, suggesting that deleting the DNAJ-interacting domain of PKL does not prevent its silencing function (Fig 2F-G). By contrast, DNAJ41-ZF108 could silence *FWA* while DNAJ41xPKL failed to do so, suggesting that PKL interaction is required for DNAJ41-mediated silencing (Fig 2F-G).

This suggests that the DNAJs are key to localization of the PKL-DNAJ-ATRX complex, but are not themselves capable of silencing without PKL.

### DNAJs recruit PKL to targeted genes via a group of transcription factors

The DNAJs do not contain well-characterized ‘reader’ domains that explain their recruitment to chromatin. Therefore, we investigated the mechanism by which DNAJs guide PKL-DNAJ-ATRX complex recruitment further. Using the IP-MS data from all three DNAJs, we identified a group of transcription factors and their associated complexes that repeatedly appeared as interactors (Fig 3A-B, Fig S4A). These interactors included WRKY transcription factor WRKY39 and its partners, the four OBERON (OBE1-4) proteins, and DE1 Binding Factor 1 (DF1) (Fig 3A-B, Fig S4A). Thus, it is possible that these transcription factors recruit DNAJs. Since four OBE proteins were significantly enriched in the DNAJ41 IP-MS, and OBE1 ChIP-seq has been published, we plotted OBE1 binding by ChIP-seq over our PKL peaks and found that OBE1 is enriched over PKL-DNAJ-ATRX complex targets (Fig 3C) [44]. This raised the possibility that DNAJs may recruit the PKL-DNAJ-ATRX complex via interactions with these transcription factors.

**Figure 3.**
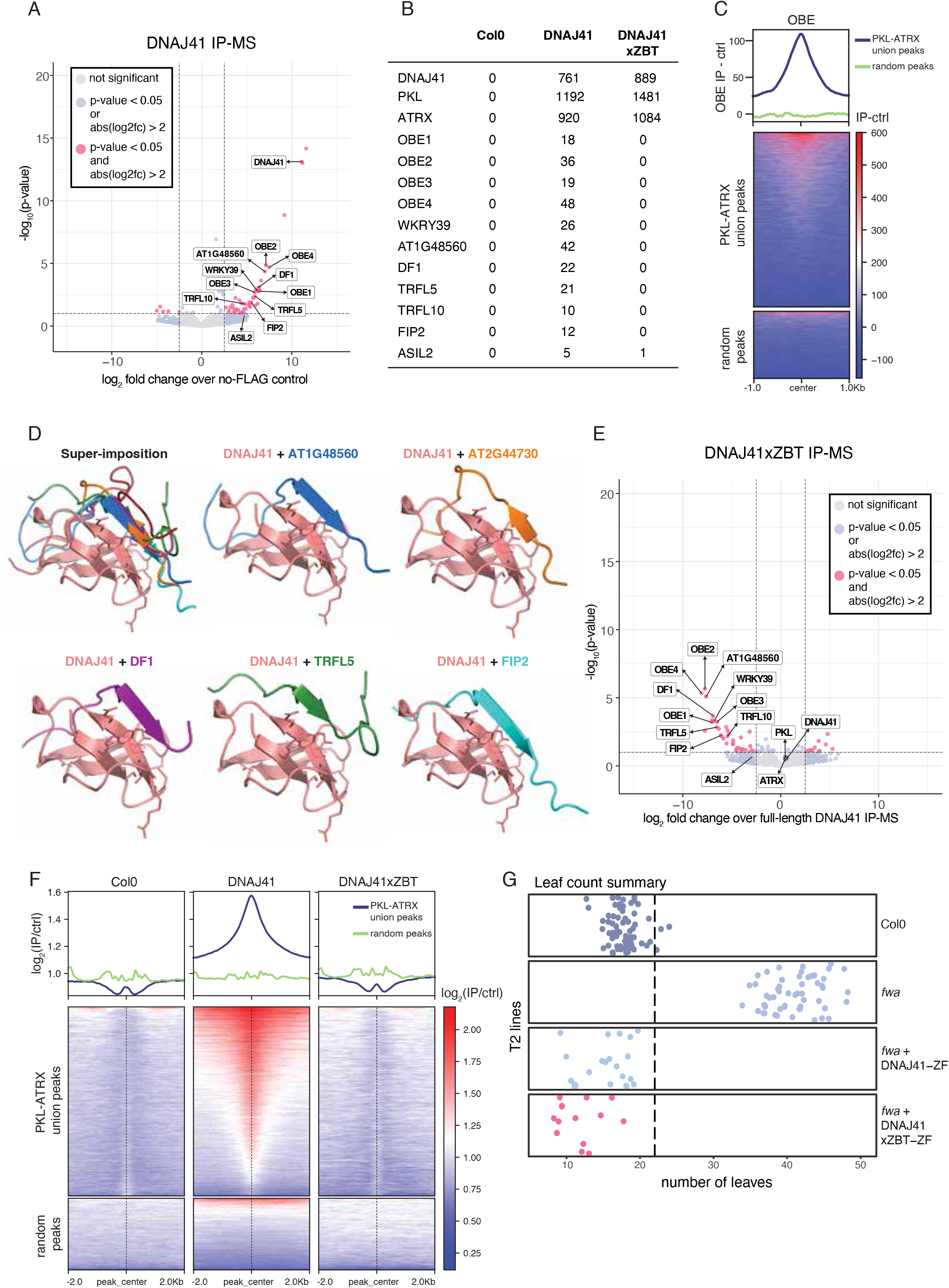
DNAJs recruit PKL to targeted genes via a group of TFs A. Volcano plot showing protein enrichment from DNAJ41 IP-MS. Pink dots represent significantly enriched or depleted proteins compared to Col-0 control. **B**. Normalized protein enrichment summary of Col-0, DNAJ41, and DNAJ41xZBT. **C**. Metaplot and heatmap showing the input normalized ChIP-seq signal of OBE1 at PKL-DNAJ-ATRX complex peaks and random peaks. **D**. AF3 modeling of the interacting interfaces between DNAJ41 and five transcription factors. **E**. Volcano plot showing normalized protein enrichment from DNAJ41xZBT IP-MS normalized to DNAJ41 IP-MS. **F**. ChIP-seq enrichment of Col-0, DNAJ41, and DNAJ41xZBT at PKL-DNAJ-ATRX complex peaks vs. random peaks. **G**. Leaf count at bolting of Col-0, *fwa*, and T2 lines of DNAJ41-ZF108 and DNAJ41xZBT-ZF108.

To test this hypothesis, we investigated how DNAJ41 interacts with these transcription factors. From AF3 modeling, we found that a region in the N-terminal domain of DNAJ41 is predicted to form close contacts with short linear motifs (SLiMs) buried in the IDR regions of each of these transcription factors (Fig 3D; Fig S4B-H, Table S3-12). This domain appears to form a C4 zinc-binding motif, with four cysteine residues coordinating a single zinc atom (Fig S4I). Based on this feature, we refer to it as the zinc-binding Tudor-like domain (ZBT). The short proline-containing motifs in the SLiMs are predicted to fold into a beta sheet in the presence of the beta-barrel of the ZBT domain (Fig 3D), and the proline inserts into a pocket and interacts with aromatic residues of the ZBT domain (Fig 3D, Table S3-12). We therefore asked whether deleting the ZBT domain of DNAJ41 would abolish its interactions with these transcription factors, disrupting DNAJ41 binding to chromatin. We created a transgene with FLAG-tagged DNAJ41 lacking the ZBT domain (DNAJ41xZBT), verified DNAJ41xZBT protein expression (Fig S2E), and performed IP-MS.

DNAJ41xZBT-FLAG lost almost all interaction with the transcription factors enriched in DNAJ41 IP-MS but still interacted with PKL and ATRX (Fig 3B and 3E, Fig S4A). Next, we performed ChIP-seq to assess whether removing the ZBT domain alters DNAJ41 binding to chromatin. DNAJ41xZBT was strongly depleted from its originally targeted regions (Fig 3F). To determine whether this loss of binding was due to loss of TF interaction or due to altered DNAJ41xZBT function or stability, we again introduced DNAJ41xZBT-ZF108 into the *fwa* epiallele mutant to test whether DNAJ41xZBT could silence the *FWA* gene. Like DNAJ41-ZF108, DNAJ41xZBT-ZF108 was still capable of silencing *FWA* (Fig 3G, Fig S4J). Overall, our results provide a mechanism for PKL recruitment to chromatin bound by specific TFs, with the DNAJs acting as a bridge between these two components.

### PKL is required for deposition of H3.3 to attenuate chromatin opening to tune gene expression

PKL has been reported to have widely varying effects on transcription, in some cases promoting silencing while in other cases promoting gene activation [19, 39]. The mechanism by which PKL regulates gene expression remains unclear. Since our PKL IP-MS results showed strong enrichment for ATRX (Fig 1A, 1C, and 1E), an H3.3 chaperone, we hypothesized that the PKL-DNAJ-ATRX complex regulates transcription in part by modulating H3 variant composition. At nucleosome-dense regions, PKL and ATRX may promote H3.3-enrichment relative to H3.1, thereby maintaining a local H3.3/H3.1 balance that supports a more transcriptionally permissive state. In contrast, at nucleosome-depleted regions, PKL-dependent H3.3 deposition may exert the opposite effect by promoting higher nucleosome density and thus a more closed chromatin state, in part via H3K27me3 spreading [45–47]. To investigate this, we generated transgenic lines expressing FLAG-tagged H3.3 or H3.1 in wild-type Columbia-0 (Col-0), *pkl*, and *atrx* mutants and performed ChIP-seq. Loss of PKL altered the positioning of H3.3 relative to PKL-DNAJ-ATRX complex binding sites, with loss of H3.3 at the original peak centers and a shift to adjacent regions (Fig. 4A, Fig S5A-C). We then examined H3.3 distribution per locus and found that the H3.3 center-to-flank ratio shows a significant decrease upon PKL loss (Fig. S5C). These observations suggest that PKL moves H3.3-containing nucleosomes closer to PKL-DNAJ-ATRX complex binding sites. By contrast, *atrx* mutants showed a significant reduction in H3.3 levels at targeted regions without a similar lateral shift in positioning (Fig. 4B), consistent with ATRX’s reported role in assembling H3.3-containing nucleosomes. Interestingly, the degree of H3.3 reduction observed in *atrx* mutants was much stronger than in *pkl* mutants, indicating that ATRX also deposits H3.3 independently of PKL. H3.1 was depleted at PKL-DNAJ-ATRX complex peaks but modestly increased in both *pkl* and *atrx* mutants (Fig. 4C-D), suggesting that PKL and ATRX both function to antagonize H3.1 accumulation at these sites. Correlation analysis further demonstrated that regions highly enriched for both PKL and ATRX exhibited substantial decreases in H3.3 and concurrent increases in H3.1 upon loss of PKL and ATRX (Fig. 4E-H).

**Figure 4.**
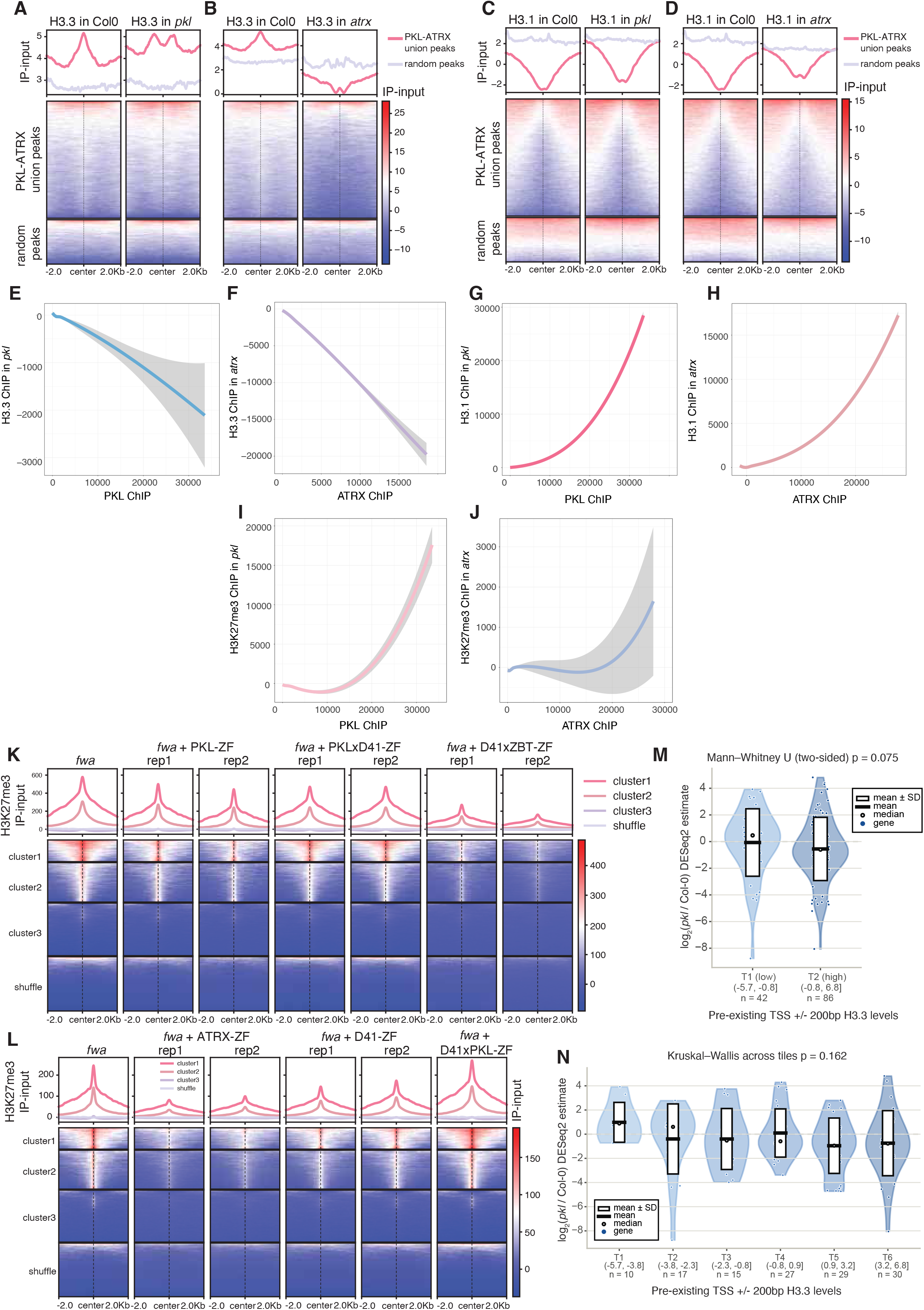
PKL together with ATRX deposits H3.3 while preventing H3.1 accumulation at targeted regions. A-B. H3.3 enrichment in **A**. Col-0, *pkl*, and **B**. *atrx* over PKL-DNAJ-ATRX complex peaks and random peaks. **C-D**. H3.1 enrichment in **C**. Col-0, *pkl*, and **D**. *atrx* over PKL-DNAJ-ATRX complex peaks and random peaks. **E-F**. Regression analysis comparing H3.3 changes and **E**. PKL or **F**. ATRX enrichment. **G-H**. Same as E-F but comparing H3.1 changes and **G**. PKL or **H**. ATRX enrichment. **I-J**. Same as E-F but comparing H3K27me3 changes and **I**. PKL or **J**. ATRX enrichment. **K-L**. Metaplots and heatmaps showing H3K27me3 ChIP-seq signal normalized to input from *fwa* and corresponding T2 lines of ZF108 transgenic plants at ZF108 off-target peaks. ZF108 peaks are clustered according to the wild-type H3K27me3 levels of *fwa*. **K** and **L** represent two independent batches of plants and experiments. Line colors in L same as in K. **M-N**. RNA-seq log2 fold change (*pkl/*wild-type) of *pkl* DEGs that show H3.3 loss at TSSs. *pkl* DEGs (n=128) are binned into **M**. 2-tiles or **N**. 6-tiles based on the pre-existing H3.3 levels at the TSS of the gene in Col-0, with a cutoff of p-value <0.05. Box indicates mean +/−SD, median is indicated by black dot, and blue points indicate individual genes.

Because H3.1 is preferentially associated with repressive chromatin states, including H3K27me3-marked regions [45–47], we examined how PKL-dependent histone variant changes affected H3K27me3 levels. We found that H3K27me3 levels increased at PKL-targeted regions upon PKL loss, correlating with increased H3.1 accumulation (Fig. 4G,I; Fig S5D). The increase in H3K27me3 levels primarily flanks ATRX binding sites in the *atrx* mutant (Fig. 4J; Fig. S5D). We reasoned that the distinct patterns of H3K27me3 accumulation in the *pkl* and *atrx* mutants may reflect differences in the H3.1/H3.3 ratio at the peak centers. These findings further suggest that ATRX may act redundantly with other histone chaperones to limit H3.1 accumulation at sites bound by the PKL complex. We next asked whether changes in H3.1 and H3.3 in the *pkl* mutant are associated with the redistribution of H3K27me3 (Fig. S5E–G). At peaks that lose H3.3 in *pkl*, increased H3.1 occupancy coincides with H3K27me3 gain (Fig. S5E). To assess which histone variant features best explain H3K27me3 redistribution, we trained machine learning models to predict H3K27me3 changes from H3.1 and H3.3 levels in WT and their changes upon loss of PKL. These models identified H3.1 levels in *pkl* as the strongest predictor of H3K27me3 redistribution (Fig. S5F-G). Simultaneously, through the gain-of-function approach, we also observed that recruitment of ZF108-tagged PKL, ATRX, and DNAJ41, but not DNAJ41xPKL constructs, resulted in a significant reduction of H3K27me3 at the *FWA* promoter and at hundreds of off-target sites where ZF-108 is also known to bind [42](Fig. 4K-L). At *FWA* loci, where ZF108-tagged PKL, ATRX, and DNAJ41 cause transcriptional silencing, we observe an increase in total H3 levels at the previously nucleosome-depleted *FWA* promoter, indicative of H3.3 accumulation (Fig S5H). These results suggest a model in which the PKL-DNAJ-ATRX complex fine-tunes the positioning of H3.3, reducing H3.1 and preventing the accumulation of H3K27me3. However, mass spectrometry studies showed that both H3.1 and H3.3 can be trimethylated on lysine 27 [47]. This suggests that the PKL complex may act primarily through H3.1 to antagonize H3K27me3 accumulation. Therefore, our results support a model in which changes in H3.1 influence the distribution of H3K27me3 at PKL binding sites.

To examine how changes in H3.3 localization might influence transcriptional output, we examined genes that showed significant expression changes in the *pkl* mutant and analyzed how pre-existing histone levels correlated with their transcriptional changes upon PKL loss. We found that for genes with relatively high H3.3 levels at transcription start sites (TSSs) in wild type, loss of PKL causes an average mild downregulation compared to wild-type (Fig 4M). High H3.3 occupancy predicts the reduction in gene expression in the *pkl* mutant when we further stratified genes into 6 tiles based on the pre-existing H3.3 level (Fig. 4N). However, due to the limited number of *pkl* DEGs, this relation is not significant (Fig. 4N). This slight correlation between pre-existing H3.3 enrichment at TSSs and gene downregulation is absent when we performed the same analysis on *pkl* DEGs without H3.3 loss (Fig S6A-B). These results suggest that the transcriptional outcome of PKL-mediated changes in H3.3 depends on the pre-existing histone landscape.

## Discussion

Our findings define a novel PKL-DNAJ-ATRX complex, revealing that a novel group of DNAJ proteins guides the recruitment of the complex to target genes. While IDRs are widely recognized for their role in phase-separation-mediated transcriptional control [48, 49], they are also known to contain interaction modules or scaffolds that help assemble chromatin complexes and tune transcriptional outcomes [31, 50, 51]. We show that a specific set of transcription factors has SLiMs in their IDRs that interact with the ZBT domain in the DNAJs to recruit the PKL-DNAJ-ATRX complex. The ZBT domain described here is present in many additional DNAJ homologs in *Arabidopsis*, including in the protein SILENZIO, where it likely mediates the interaction with a similar SLIM domain of its binding partner ACD21 [52]. Therefore, the ZBT domain in other uncharacterized DNAJ proteins is also likely to mediate additional important SLIM interactions. In addition, we showed that an α-helix in DNAJ interacts with the SANT–SLIDE domain of PKL, thereby facilitating recruitment of the PKL–DNAJ–ATRX complex to chromatin and its regulation of gene expression. Together, these findings indicate that coordinated protein–protein interactions involving both ordered and disordered domains are important for transcriptional regulation by this complex.

However, we cannot rule out the possibility that deleting the DNAJ α-helix may also disrupt other, as-yet-uncharacterized functions of the protein. Further studies will be required to distinguish these possibilities.

The loss of PKL has been associated with both gene activation and silencing, but the underlying mechanism has remained unclear. Our study provides a potential mechanistic explanation by showing that PKL, in concert with ATRX, deposits H3.3 at target sites and also can prevent the accumulation of H3K27me3. In the *pkl* mutant, this can lead to different effects on gene expression at H3.3-rich vs. H3.3-poor regions. It therefore appears that changes in H3.3 occupancy, together with the local chromatin context at PKL-targeted sites, collectively determine the transcriptional impact of PKL. However, whether PKL-mediated H3.3 changes are dependent merely on ATRX remains unclear, and further characterization incorporating both loss-of-function and gain-of-function approaches is needed to address this.

Establishing a functional coordination between a CHD3 homolog (PKL) and ATRX to drive H3.3 deposition diverges significantly from the canonical animal models. In animals, CHD3/CHD4 proteins function primarily within the NuRD complex, whereas ATRX acts with the H3.3 chaperone DAXX to deposit H3.3 at telomeric, pericentromeric, and other heterochromatic regions [2, 53–56]. Plants thus appear to have evolved a unique link between these different epigenetic mechanisms.

In summary, this work uncovers a new mechanism for CHD3 chromatin remodeler recruitment by DNAJ proteins that functions through H3.3 deposition, furthering our understanding of epigenetic gene regulation in plant systems.

## Methods and Material

### Plant Materials and Growth Conditions

*Arabidopsis thaliana* plants (Col-0 ecotype) were grown under long-day conditions (16 h light/8 h dark). Seedlings of Col-0 and *pkl* mutants were harvested after 10 days of incubation under long-day conditions. The T-DNA insertion lines used in this study include pkl (GABI_273E06) and atrx (SALK_025687.49.00.x). Transgenic plants were produced via floral dipping with Agrobacterium tumefaciens (AGL0 strain).

### Plasmid Construction and Plant Transformation

FLAG-tagged proteins for ChIP-seq experiments were generated using the Gateway-compatible binary destination vector pEG302-effector (gDNA)-3xFLAG. Genomic sequences spanning from the native promoter (~1.5 kb upstream of the 5’ UTR or intergenic regions) to the end of the endogenous gene (excluding the stop codon) were cloned into pENTR D-TOPO vectors (Invitrogen) and transferred into the destination vector via Gateway LR Clonase II (Invitrogen). For ZF108 targeting experiments, pEG302-effector (gDNA)-3xFLAG-ZF108 was used, which includes a ZF108 motif alongside the FLAG tag.

Plasmids were introduced into Col-0 (FLAG-tagged DNAJ40, DNAJ41, DNAJ68, PKL, ATRX, and their truncated versions) or *fwa* (ZF targeting experiments) using the floral dip method. After hygromycin (FLAG constructs) or Basta (ZF constructs) selection of transgene-positive plants (T1 plants), we verified protein expression by western blot, then propagated two independent lines with similar protein levels to the next generation (T2) for IPMS and ChIP-seq experiments.

### IP-MS

Ten grams of unopened floral buds from Col-0 wild-type and FLAG-tagged transgenic plants were ground into fine powder in liquid nitrogen. Samples were resuspended in 25 mL IP buffer (50 mM Tris-HCl pH 8.0, 150 mM NaCl, 5 mM EDTA, 10% glycerol, 0.1% Tergitol) and homogenized until lump-free using a Dounce homogenizer. Lysates were filtered through Miracloth and incubated with 250 μL anti-FLAG M2 magnetic beads (Sigma) at 4°C for 2 hours. Beads were washed with IP buffer and eluted with TBS containing 250 μg/mL 3xFLAG peptides. Eluted proteins were precipitated with trichloroacetic acid (Sigma) and subjected to MS analysis as described previously[57].

### Flowering time measurement

T1 transgenic plants (first generation after dipping) expressing DNAJ41-ZF108, PKL-ZF108, ATRX-ZF108, or their respective truncated versions were selected after transgene expression was verified by western blot. Flowering time was evaluated in the following (T2) generation across multiple independent lines.

Seeds collected from the selected T1 lines were screened using Basta to confirm the presence of the transgene. After bolting, the numbers of rosette and cauline leaves were counted for each plant. Each point in the dot plots in Fig. 2 represents the total leaf count of an individual plant. The cutoff line indicates plants with 22 or fewer leaves.

### RT-qPCR

RNA was extracted from rosette leaves of 3–4-week-old plants using the Zymo Direct-zol RNA MiniPrep kit. 400 ng to 1 μg of total RNA was reverse-transcribed using Superscript III First Strand Synthesis Supermix (Invitrogen). qPCR was performed using iQ SYBR Green Supermix (Bio-Rad), with FWA expression normalized to IPP2 (ISOPENTENYL PYROPHOSPHATE DIMETHYLALLYL PYROPHOSPHATE ISOMERASE 2). Primer sequences are listed below.

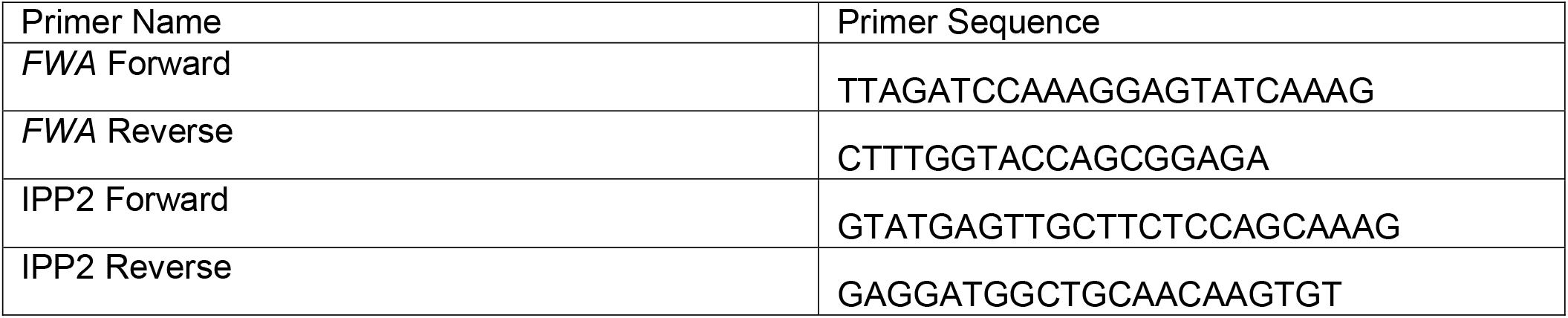

### ChIP-seq

All buffers used for ChIP-seq experiments were supplemented with PMSF, Benzamidine, and cOmplete™ Protease Inhibitor Cocktail (Sigma). Unopened flower buds (~1–2 g) or rosette leaves (~2–4 g) from T2 lines were ground in liquid nitrogen and dissolved in nuclei isolation buffer (50 mM HEPES, 1 M sucrose, 5 mM KCl, 5 mM MgCl2, 0.6% Triton X-100). Crosslinking was performed by adding formaldehyde to a final concentration of 1% followed by incubation for 12 minutes, then quenched with 125mM Glycine.

Nuclei were sequentially washed with extraction buffers containing sucrose gradients before lysis in nuclei lysis buffer (50 mM Tris-HCl pH 8, 10 mM EDTA, 1% SDS). Chromatin was sheared using a Bioruptor Plus sonicator (30 seconds ON/30 seconds OFF, High setting, 22 cycles). Immunoprecipitation was performed overnight at 4°C using anti-FLAG antibody (Sigma), followed by incubation with Protein A/G Dynabeads for 2 hours. Samples were reverse crosslinked overnight at 65°C in NaCl-containing elution buffer before protein digestion with Proteinase K and purification via Phenol:Chloroform extraction. Libraries were prepared using Ovation Ultra Low System V2 (NuGEN) and sequenced on Illumina NovaSeq.

### RNA-seq

Biological triplicates were generated for each genotype. Inflorescences containing unopened flower buds from 5–6-week-old plants or rosette leaves from 4–5-week-old plants were collected as biological replicates and frozen in liquid nitrogen. RNA was extracted using the Zymo Direct-zol RNA MiniPrep kit. RNA-seq libraries were prepared using TruSeq Stranded mRNA kits (Illumina), with sequencing performed on Illumina NovaSeq.

### ChIP-seq analysis

Low-quality read ends and Illumina adapters were removed using Trim Galore (v0.6.7, Babraham Institute). Reads were aligned to the Arabidopsis thaliana reference genome (TAIR10: https://www.arabidopsis.org/index.jsp) using Bowtie2 (v2.3.4) [58], allowing only uniquely mapped reads. PCR duplicates were removed using MarkDuplicates.jar from the Picard tools suite (v3.1.0, Broad Institute), and BAM file indexes were generated with Samtools (v1.9). BigWig files for visualization were created using deeptools (v3.0.2) bamCoverage with the options --normalizeUsing RPGC and --binSize 10 [59]. For correlation analysis between ChIP-seq signals and mCG density, samples were normalized to the no-FLAG control using deeptools bamCompare with the options --scaleFactorsMethod readCount, --binSize 10, and --operation log2. Data visualization was performed in R using ggplot with the geom_smooth option. ChIP-seq peaks were called using MACS2 (v2.1.1) with an FDR cutoff of 0.05 [60]. Peaks detected in anti-FLAG Col-0 controls were removed from the peak files.

### RNA-seq analysis

RNA-seq reads were trimmed Trim Galore, as above. The filtered reads were then aligned to TAIR10 using STAR (v2.7.11a) [61]. Only uniquely mapped reads with less than 5% mismatches were retained. BigWig files were generated using deeptools (v3.0.2) bamCoverage with the options --normalizeUsing RPGC and --binSize 10 [59]. To quantify expression, HTSeq (v0.13.5) was used to obtain read counts over genes/TEs [62]. Differential expression analysis was conducted using DESeq2 (v1.42.0), with a cutoff of padj < 0.05 and |log2FC| ≥ 1.

### Analysis of the impact of pre-existing H3.3 occupancy on gene expression

Pre-existing H3.3 levels were quantified over the TSS +/−200bp using the Col-0 H3.3 ChIP-seq signal after subtracting input. Genes were ranked by pre-existing H3.3 and divided into eight equal-sized octiles. The lowest and highest octiles were excluded and the remaining six, covering the central 75% of genes, were relabeled T1–T6. For the two-tile analysis, the same genes were split at the genome-wide median into a low tile and a high tile, so that both analyses use identical genes and nested boundaries. The extreme octiles were excluded as the highest octile has a heavy right tail and contains the extreme outliers of the data set. Tiles were defined for all genes genome-wide.

Differential expression between *pkl* and Col-0 floral buds was analyzed with DESeq2 (see RNA-seq section above). Genes with p < 0.05 were used, without a fold-change filter, because these analyses compare the distribution of expression changes rather than calling individual differentially expressed genes, and an FDR threshold left too few genes for stratification. Of the 870 genes, 646 are present in the annotation used here, and 458 fell within the six retained octiles. Among the 458 retained pkl DEGs, 128 lost H3.3 (T1–T6: n = 10, 17, 15, 27, 29 and 30; low/high tiles: n = 42 and 86); the remaining 330 genes served as the no-loss control.

Violin plots show log2FC per tile with every gene overlaid; the box spans the mean ± SD, the thick line marks the mean and the open circle the median. Because log2FC values within tiles are heavy-tailed, group sizes are small and unequal, and variances differ between tiles, we used rank-based two-sided tests (SciPy 1.17.0). Differences among the six tiles were tested with the Kruskal–Wallis H test and the monotonic trend across tiles with Spearman’s rank correlation between tile index and log2FC; the two tiles were compared with the Mann–Whitney U test.

### Analysis of histone variant and histone modification

We carried out a two-dimensional binned summary of how H3.1 and H3.1 changes influence the gain of H3K27me3 at PKL complex peaks. Three statistics: H3.3 change, H3.1 change, and H3K27me3 change in the *pkl* mutant, were summarized over each peak. All three are base-pair-weighted means over the peak interval. Then peaks were split into six equal-sized groups by their H3.3 change, and independently into six groups by their H3.1 change. For the peaks in a cell within the heatmap, the intensity indicates the percentage whose H3K27me3 change lies above the genome-wide median.

We trained a panel of machine-learning models (scikit-learn 1.8.0) to predict the H3K27me3 change in *pkl* at PKL-complex peaks from six features (H3.3 and H3.1 in wild type, H3.3 and H3.1 in *pkl*, and the *pkl*-minus-wild-type change of each). Hyperparameters were selected by five-fold cross-validation on 80% of peaks, performance was measured on the remaining 20% and by leave-one-chromosome-out validation, and the random-forest model performed best. Permutation feature importance (mean decrease in test R^2^ over 20 random shuffles) ranked H3.1 level in *pkl* first (0.13) and the H3.1 change second (0.04), followed by the H3.3 change (0.03); permuting the three H3.1 features together reduced R^2^ by 0.32, compared with 0.06 for the three H3.3 features. The same ranking was obtained with every model and with the gain-versus-loss classifier.

## Data availability

The high-throughput sequencing data generated in this paper will be deposited in the Gene Expression Omnibus [63]. The TAIR10 genome is available at https://www.arabidopsis.org/index.jsp.

## Acknowledgements

We thank Jiamu Du for advice on modeling the ZBT domain. We thank Mahnaz Akhavan and the UCLA BSCRC BioSequencing Core for the sequencing support. This work was supported by George G. & Betsy H. Laties Graduate Fellowship in Molecular Plant Biology to S.W. S.E.J. is an investigator of the Howard Hughes Medical Institute.

## References

1. Kouzarides, T., Chromatin modifications and their function. Cell, 2007. 128(4): p. 693–705.

2. Goldberg, A.D., C.D. Allis, and E. Bernstein, Epigenetics: a landscape takes shape. Cell, 2007. 128(4): p. 635–8.

3. Law, J.A. and S.E. Jacobsen, Establishing, maintaining and modifying DNA methylation patterns in plants and animals. Nat Rev Genet, 2010. 11(3): p. 204–20.

4. Struhl, K. and E. Segal, Determinants of nucleosome positioning. Nat Struct Mol Biol, 2013. 20(3): p. 267–73.

5. Marfella, C.G. and A.N. Imbalzano, The Chd family of chromatin remodelers. Mutat Res, 2007. 618(1-2): p. 30–40.

6. Torchy, M.P., A. Hamiche, and B.P. Klaholz, Structure and function insights into the NuRD chromatin remodeling complex. Cell Mol Life Sci, 2015. 72(13): p. 2491–507.

7. Lai, A.Y. and P.A. Wade, Cancer biology and NuRD: a multifaceted chromatin remodelling complex. Nat Rev Cancer, 2011. 11(8): p. 588–96.

8. Burgold, T., et al., The Nucleosome Remodelling and Deacetylation complex suppresses transcriptional noise during lineage commitment. EMBO J, 2019. 38(12).

9. Knock, E., et al., The methyl binding domain 3/nucleosome remodelling and deacetylase complex regulates neural cell fate determination and terminal differentiation in the cerebral cortex. Neural Dev, 2015. 10: p. 13.

10. Ogas, J., et al., Cellular differentiation regulated by gibberellin in the Arabidopsis thaliana pickle mutant. Science, 1997. 277(5322): p. 91–4.

11. Ogas, J., et al., PICKLE is a CHD3 chromatin-remodeling factor that regulates the transition from embryonic to vegetative development in Arabidopsis. Proc Natl Acad Sci U S A, 1999. 96(24): p. 13839–44.

12. Aichinger, E., et al., The CHD3 chromatin remodeler PICKLE and polycomb group proteins antagonistically regulate meristem activity in the Arabidopsis root. Plant Cell, 2011. 23(3): p. 1047–60.

13. Park, J., et al., Gibberellin Signaling Requires Chromatin Remodeler PICKLE to Promote Vegetative Growth and Phase Transitions. Plant Physiol, 2017. 173(2): p. 1463–1474.

14. Xu, M., et al., Epigenetic Regulation of Vegetative Phase Change in Arabidopsis. Plant Cell, 2016. 28(1): p. 28–41.

15. Liang, Z., et al., The transcriptional repressors VAL1 and VAL2 mediate genome-wide recruitment of the CHD3 chromatin remodeler PICKLE in Arabidopsis. Plant Cell, 2022. 34(10): p. 3915–3935.

16. Song, M., et al., PKL mediates H3K4me2 modification and spatial gene congregation in chromatin regulation. Nucleic Acids Res, 2025. 53(22).

17. Jing, Y., et al., Arabidopsis chromatin remodeling factor PICKLE interacts with transcription factor HY5 to regulate hypocotyl cell elongation. Plant Cell, 2013. 25(1): p. 242–56.

18. Aichinger, E., et al., CHD3 proteins and polycomb group proteins antagonistically determine cell identity in Arabidopsis. PLoS Genet, 2009. 5(8): p. e1000605.

19. Liang, Z., et al., PICKLE-mediated nucleosome condensing drives H3K27me3 spreading for the inheritance of Polycomb memory during differentiation. Mol Cell, 2024. 84(18): p. 3438–3454 e8.

20. Verma, A.K., et al., The expanding world of plant J-domain proteins. CRC Crit Rev Plant Sci, 2019. 38(5-6): p. 382–400.

21. Pulido, P. and D. Leister, Novel DNAJ-related proteins in Arabidopsis thaliana. New Phytol, 2018. 217(2): p. 480–490.

22. Qiu, X.B., et al., The diversity of the DnaJ/Hsp40 family, the crucial partners for Hsp70 chaperones. Cell Mol Life Sci, 2006. 63(22): p. 2560–70.

23. Hennessy, F., et al., Not all J domains are created equal: implications for the specificity of Hsp40-Hsp70 interactions. Protein Sci, 2005. 14(7): p. 1697–709.

24. Rajan, V.B. and P. D’Silva, Arabidopsis thaliana J-class heat shock proteins: cellular stress sensors. Funct Integr Genomics, 2009. 9(4): p. 433–46.

25. Ajit Tamadaddi, C. and C. Sahi, J domain independent functions of J proteins. Cell Stress Chaperones, 2016. 21(4): p. 563–70.

26. Kampinga, H.H., et al., Function, evolution, and structure of J-domain proteins. Cell Stress Chaperones, 2019. 24(1): p. 7–15.

27. Harris, C.J., et al., A DNA methylation reader complex that enhances gene transcription. Science, 2018. 362(6419): p. 1182–1186.

28. Ichino, L., et al., MBD5 and MBD6 couple DNA methylation to gene silencing through the J-domain protein SILENZIO. Science, 2021.

29. Li, S., et al., SUVH1, a Su(var)3-9 family member, promotes the expression of genes targeted by DNA methylation. Nucleic Acids Res, 2016. 44(2): p. 608–20.

30. Zhao, Q.Q., et al., A methylated-DNA-binding complex required for plant development mediates transcriptional activation of promoter methylated genes. J Integr Plant Biol, 2019. 61(2): p. 120–139.

31. Patil, A., et al., A disordered region controls cBAF activity via condensation and partner recruitment. Cell, 2023. 186(22): p. 4936–4955 e26.

32. Li, B., M. Carey, and J.L. Workman, The role of chromatin during transcription.Cell, 2007. 128(4): p. 707–19.

33. Petty, E. and L. Pillus, Balancing chromatin remodeling and histone modifications in transcription. Trends Genet, 2013. 29(11): p. 621–9.

34. Duc, C., et al., Arabidopsis ATRX Modulates H3.3 Occupancy and Fine-Tunes Gene Expression. Plant Cell, 2017. 29(7): p. 1773–1793.

35. Wang, H., et al., LHP1 Interacts with ATRX through Plant-Specific Domains at Specific Loci Targeted by PRC2. Mol Plant, 2018. 11(8): p. 1038–1052.

36. Nie, X., et al., The HIRA complex that deposits the histone H3.3 is conserved in Arabidopsis and facilitates transcriptional dynamics. Biol Open, 2014. 3(9): p. 794–802.

37. Tanaka, M., A. Kikuchi, and H. Kamada, The Arabidopsis histone deacetylases HDA6 and HDA19 contribute to the repression of embryonic properties after germination. Plant Physiol, 2008. 146(1): p. 149–61.

38. Carter, B., et al., The Chromatin Remodelers PKL and PIE1 Act in an Epigenetic Pathway That Determines H3K27me3 Homeostasis in Arabidopsis. Plant Cell, 2018. 30(6): p. 1337–1352.

39. Jing, Y., Q. Guo, and R. Lin, The Chromatin-Remodeling Factor PICKLE Antagonizes Polycomb Repression of FT to Promote Flowering. Plant Physiol, 2019. 181(2): p. 656–668.

40. Zhang, D., et al., The Chromatin-Remodeling Factor PICKLE Integrates Brassinosteroid and Gibberellin Signaling during Skotomorphogenic Growth in Arabidopsis. Plant Cell, 2014. 26(6): p. 2472–2485.

41. Johnson, L.M., et al., SRA- and SET-domain-containing proteins link RNA polymerase V occupancy to DNA methylation. Nature, 2014. 507(7490): p. 124–128.

42. Gallego-Bartolome, J., et al., Co-targeting RNA Polymerases IV and V Promotes Efficient De Novo DNA Methylation in Arabidopsis. Cell, 2019. 176(5): p. 1068–1082 e19.

43. Soppe, W.J., et al., The late flowering phenotype of fwa mutants is caused by gain-of-function epigenetic alleles of a homeodomain gene. Mol Cell, 2000. 6(4): p. 791–802.

44. Du, P., et al., WRKY transcription factors and OBERON histone-binding proteins form complexes to balance plant growth and stress tolerance. EMBO J, 2023. 42(19): p. e113639.

45. Jiang, D. and F. Berger, DNA replication-coupled histone modification maintains Polycomb gene silencing in plants. Science, 2017. 357(6356): p. 1146–1149.

46. Stroud, H., et al., Genome-wide analysis of histone H3.1 and H3.3 variants in Arabidopsis thaliana. Proc Natl Acad Sci U S A, 2012. 109(14): p. 5370–5.

47. Jamge, B., et al., Histone variants shape chromatin states in Arabidopsis. Elife, 2023. 12.

48. Sabari, B.R., et al., Coactivator condensation at super-enhancers links phase separation and gene control. Science, 2018. 361(6400).

49. Borcherds, W., et al., How do intrinsically disordered protein regions encode a driving force for liquid-liquid phase separation? Curr Opin Struct Biol, 2021. 67: p. 41–50.

50. Shang, B., C. Li, and X. Zhang, How intrinsically disordered proteins order plant gene silencing. Trends Genet, 2024. 40(3): p. 260–275.

51. Tabar, M.S., et al., Intrinsically Disordered Regions Define Unique Protein Interaction Networks in CHD Family Remodelers. FASEB J, 2025. 39(10): p. e70632.

52. Boone, B.A., et al., ACD15, ACD21, and SLN regulate the accumulation and mobility of MBD6 to silence genes and transposable elements. Sci Adv, 2023. 9(46): p. eadi9036.

53. Tong, J.K., et al., Chromatin deacetylation by an ATP-dependent nucleosome remodelling complex. Nature, 1998. 395(6705): p. 917–21.

54. Xue, Y., et al., NURD, a novel complex with both ATP-dependent chromatin-remodeling and histone deacetylase activities. Mol Cell, 1998. 2(6): p. 851–61.

55. Drane, P., et al., The death-associated protein DAXX is a novel histone chaperone involved in the replication-independent deposition of H3.3. Genes Dev, 2010. 24(12): p. 1253–65.

56. Lewis, P.W., et al., Daxx is an H3.3-specific histone chaperone and cooperates with ATRX in replication-independent chromatin assembly at telomeres. Proc Natl Acad Sci U S A, 2010. 107(32): p. 14075–80.

57. Wu, Z., et al., REM transcription factors and GDE1 shape the DNA methylation landscape through the recruitment of RNA polymerase IV transcription complexes. Nat Cell Biol, 2025. 27(7): p. 1136–1147.

58. Langmead, B. and S.L. Salzberg, Fast gapped-read alignment with Bowtie 2. Nat Methods, 2012. 9(4): p. 357–9.

59. Ramirez, F., et al., deepTools2: a next generation web server for deep-sequencing data analysis. Nucleic Acids Res, 2016. 44(W1): p. W160–5.

60. Zhang, Y., et al., Model-based analysis of ChIP-Seq (MACS). Genome Biol, 2008. 9(9): p. R137.

61. Dobin, A., et al., STAR: ultrafast universal RNA-seq aligner. Bioinformatics, 2013. 29(1): p. 15–21.

62. Anders, S., P.T. Pyl, and W. Huber, HTSeq--a Python framework to work with high-throughput sequencing data. Bioinformatics, 2015. 31(2): p. 166–9.

63. Jacob, Y., et al., Selective methylation of histone H3 variant H3.1 regulates heterochromatin replication. Science, 2014. 343(6176): p. 1249–53.

