## Supplementary data for "J-domain proteins bridge the PKL and ATRX complex to transcription factors to fine-tune gene expression via H3.3 deposition"

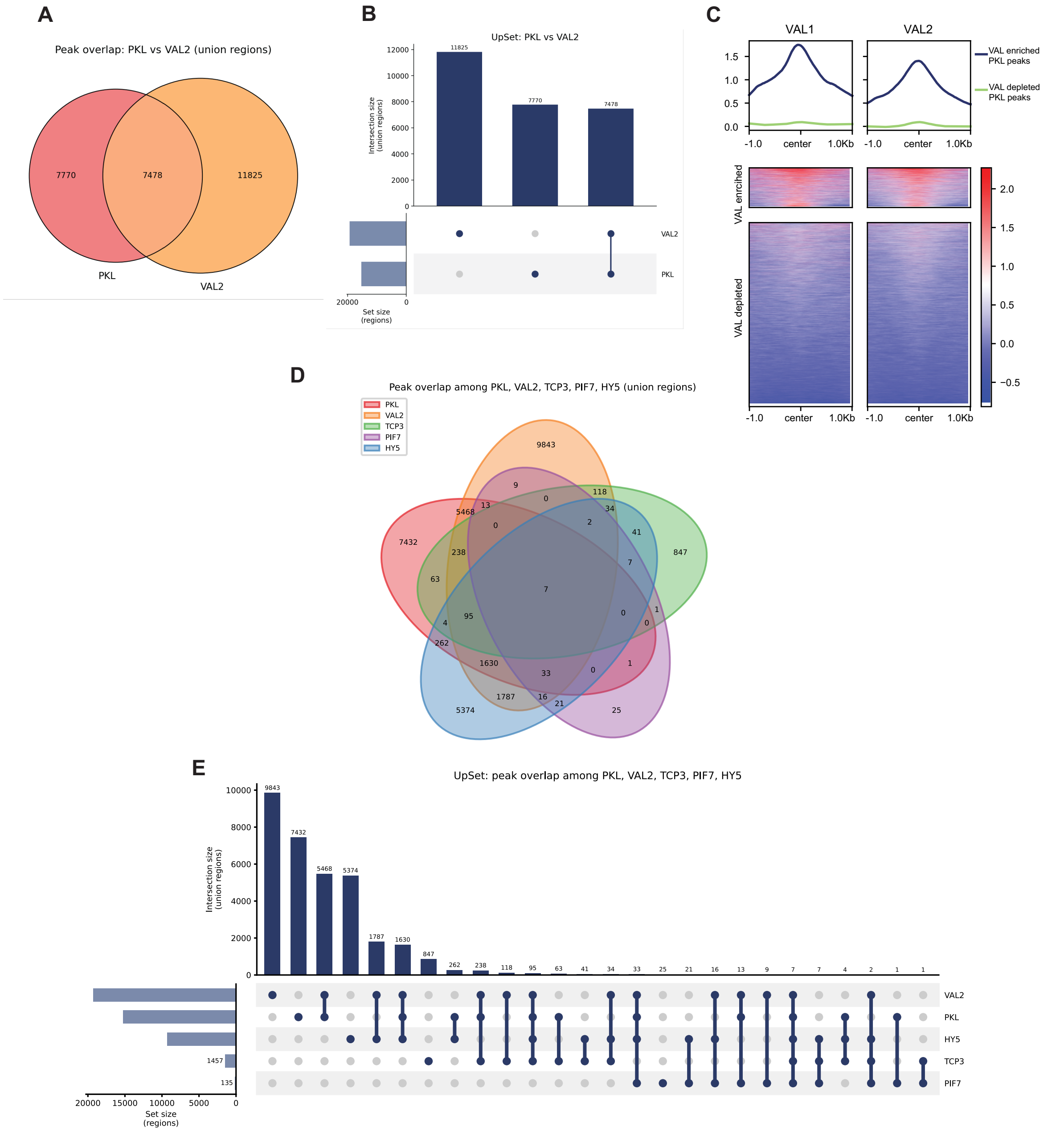


F

| | Col0 | PKL | ATRX | DNAJ40 | DNAJ41 | DNAJ68 |
|---|---|---|---|---|---|---|
| | | AT2G25170 | AT1G08600 | AT5G53150 | AT2G25560 | AT2G05230 |
| **PKL** | 0 | 136 | 100 | 104 | 91 | 108 |
| **ATRX** | 0 | 109 | 154 | 43 | 113 | 36 |
| **DNAJ40** | 0 | 61 | 13 | 72 | 1 | 10 |
| **DNAJ41** | 0 | 40 | 47 | 17 | 46 | 11 |
| **DNAJ68** | 0 | 46 | 25 | 3 | 1 | 43 |

G

| | Col0 | PKL | PKLxDNAJ | DNAJ41 | DNAJ41xPKL |
|---|---|---|---|---|---|
| | | AT2G25170 | AT2G25170 | AT2G25560 | AT2G25560 |
| **PKL** | 0 | 136 | 89 | 91 | 39 |
| **ATRX** | 0 | 109 | 46 | 113 | 28 |
| **DNAJ40** | 0 | 61 | 0 | 1 | 5 |
| **DNAJ41** | 0 | 40 | 5 | 46 | 36 |
| **DNAJ68** | 0 | 562 | 0 | 1 | 12 |

**Figure S1. A subset of PKL peaks overlaps with transcription factors VAL1/2, HY5, TCP3, and PIF7.**

**A.** Venn diagram showing the overlap between PKL and VAL2 ChIP-seq peaks. **B.** Upset plot showing distinct and shared peaks between PKL and VAL2. **C.** Input-normalized ChIP-seq signal of VAL1 and VAL2 at PKL peaks. Peaks are clustered based on enrichment in VAL1 and VAL2. **D.** Venn diagram showing the overlap among PKL, VAL2, TCP3, PIF7, and HY5. **E.** Upset plot showing the distinct and shared peaks among PKL, VAL2, TCP3, PIF7, and HY5. Unique peptide count summary of **F.** Col-0, PKL, ATRX, DNAJ40, DNAJ41, and DNAJ68 IP-MS and of **G.** Col-0, PKL, PKLxDNAJ, DNAJ41, and DNAJ41xPKL IP-MS.

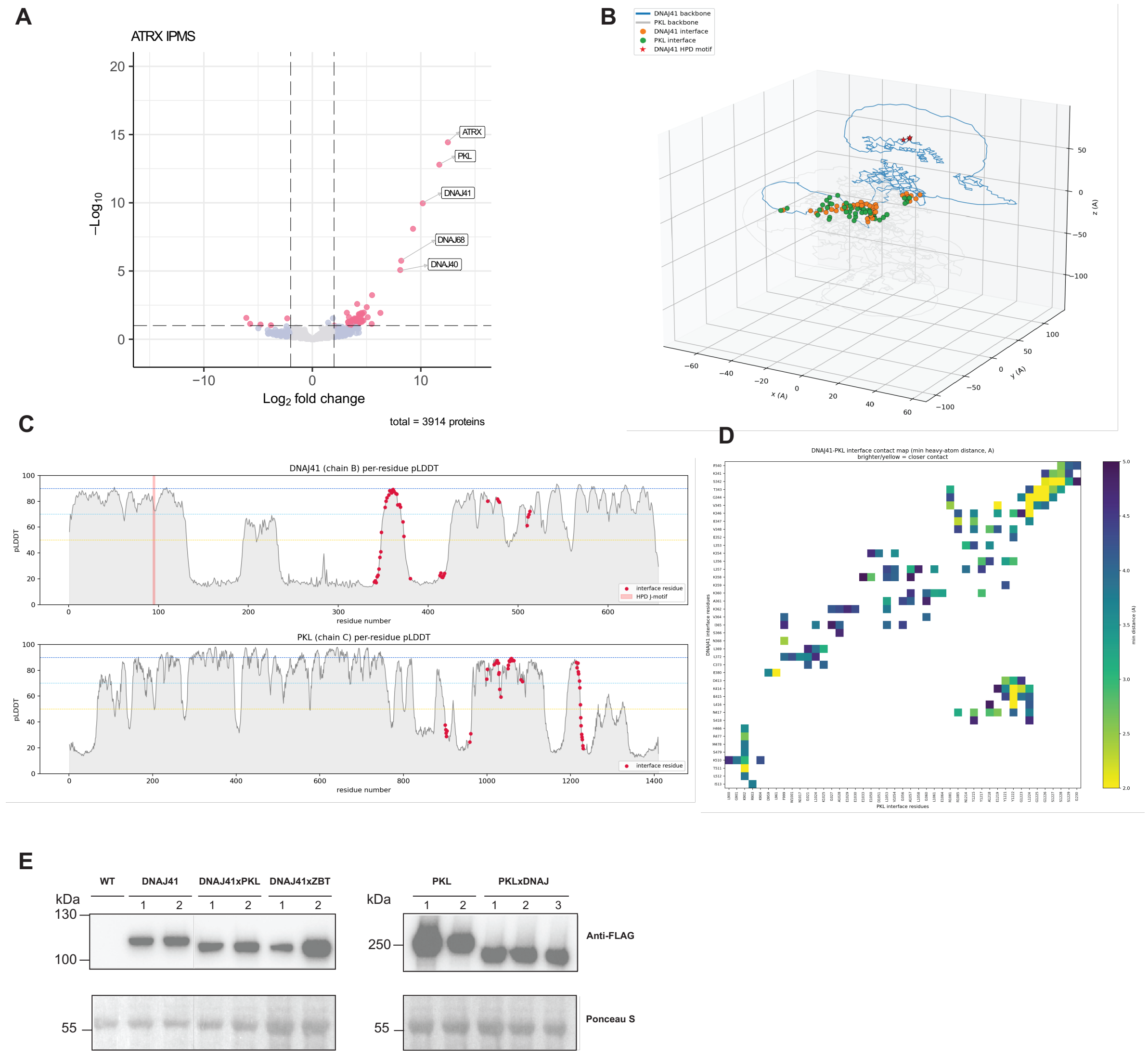


**Figure S2. DNAJs interact with PKL via the alpha-helical domain. A.** Volcano plots showing protein enrichment from ATRX IPMS. **B.** 3D contact map between DNAJ41 and PKL. **C.** pLDDT plots of DNAJ41 and PKL with interacting residues highlighted in red. **D.** 2D residue contact map between DNAJ41 and PKL. **E.** Western blots showing the protein levels of 3xFLAG-tagged DNAJ41, DNAJ41xPKL, DNAJ41xZBT, along with PKL and PKLxDNAJ. The top panel shows anti-FLAG antibody staining, while the bottom panel shows the Ponceau S staining as the loading control.

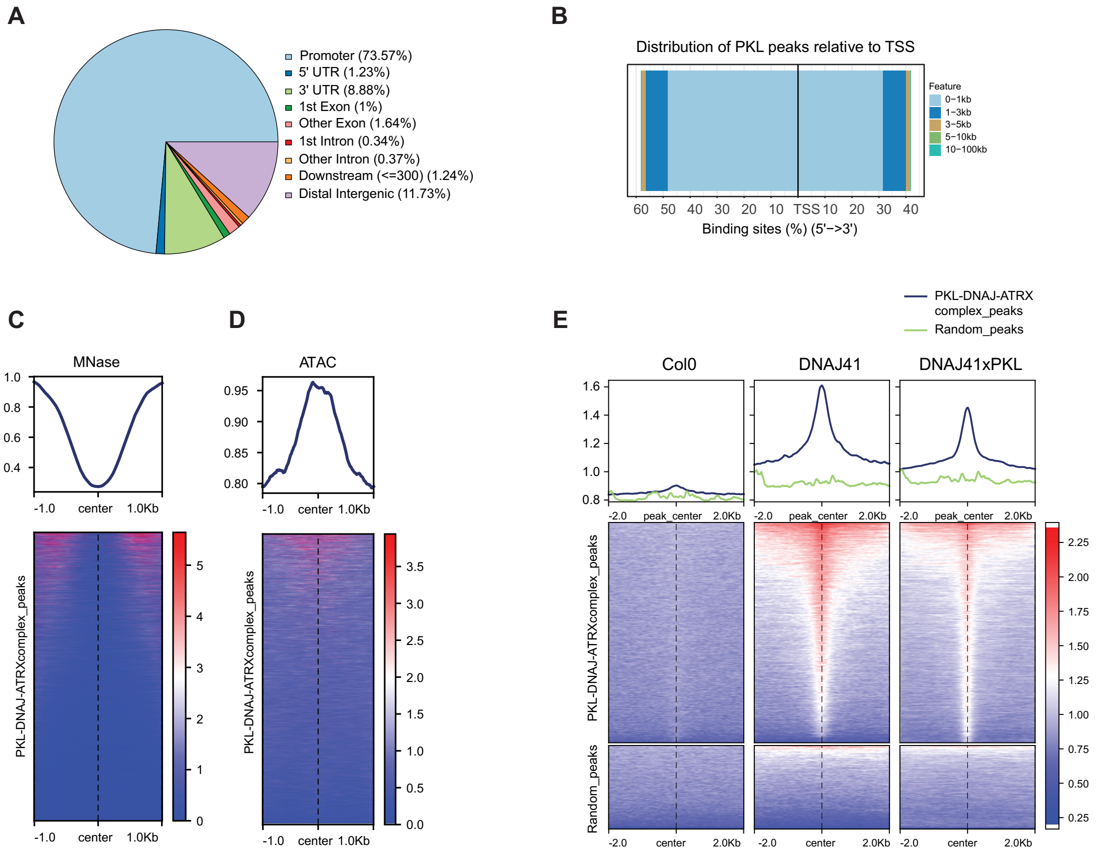


**Figure S3. PKL complex enriches at accessible gene promoters. A.** Pie chart showing the percentage of peaks associated with different gene features. **B.** Distribution of PKL peaks relative to transcription start site (TSS). PKL peaks are grouped by their relative distance to the TSS. Relative orientation of peaks to the TSS is indicated on the x-axis. Metaplots and heatmaps showing the **C.** MNase-seq and **D.** ATAC-seq signal at PKL peaks. **E.** Metaplots and heatmaps showing the ChIP-seq signal of Col-0, DNAJ41, and DNAJ41xPKL at PKL peaks and random peaks.

**A**

| | DNAJ41 | OBE1 | OBE2 | OBE3 | OBE4 | WKRY39 | AT1G48560 | DF1 | TRFL5 | TRFL10 | FIP2 | ASIL2 |
|---|---|---|---|---|---|---|---|---|---|---|---|---|
| **Col0** | 0 | 0 | 0 | 0 | 0 | 0 | 0 | 0 | 0 | 0 | 0 | 0 |
| **DNAJ41** | 46 | 9 | 17 | 15 | 28 | 9 | 22 | 7 | 8 | 6 | 7 | 5 |
| **DNAJ41xZBT** | 51 | 0 | 0 | 0 | 0 | 0 | 0 | 0 | 0 | 0 | 0 | 1 |

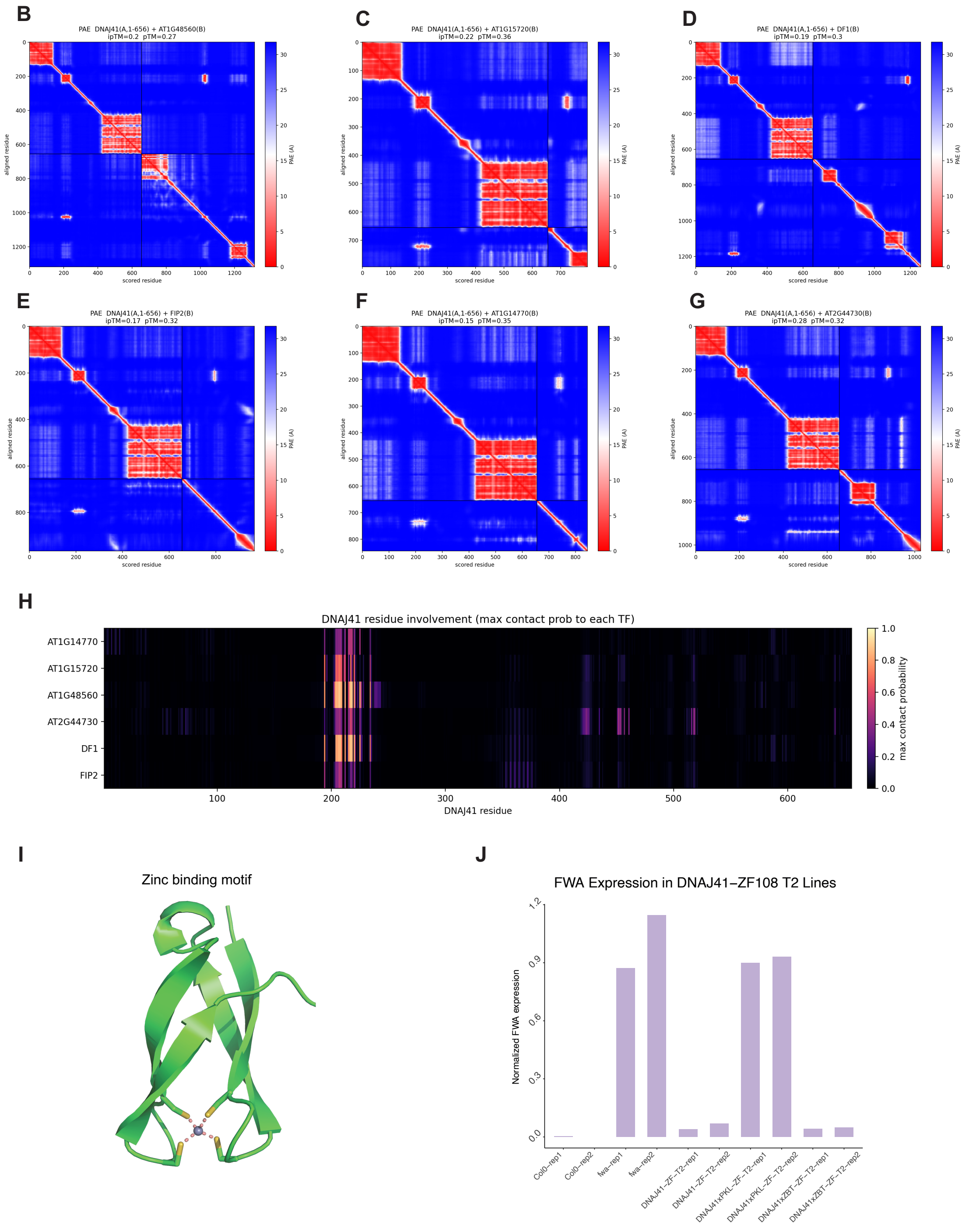

**Figure S4. AF3 predicts DNAJ41 interaction with a group of transcription factors via the ZBT domain. A.** Unique peptide count summary of Col0, DNAJ41, and DNAJ41xZBT IPMS. PAE matrices showing the interaction confidence between DNAJ41 and **B.** AT1G48560, **C.** AT1G15720, **D.** DF1, **E.** FIP2, **F.** AT1G14770, and **G.** AT2G44730. **H.** Heatmap showing the maximum contact probability of DNAJ41 residues with different transcription factors. **I.** 3D modeling of the ZBT domain with 4 cysteines binding to a zinc ion. **J.** Bar plot summarizing RT-qPCR results from the Col-0, *fwa*, and corresponding T2 lines.

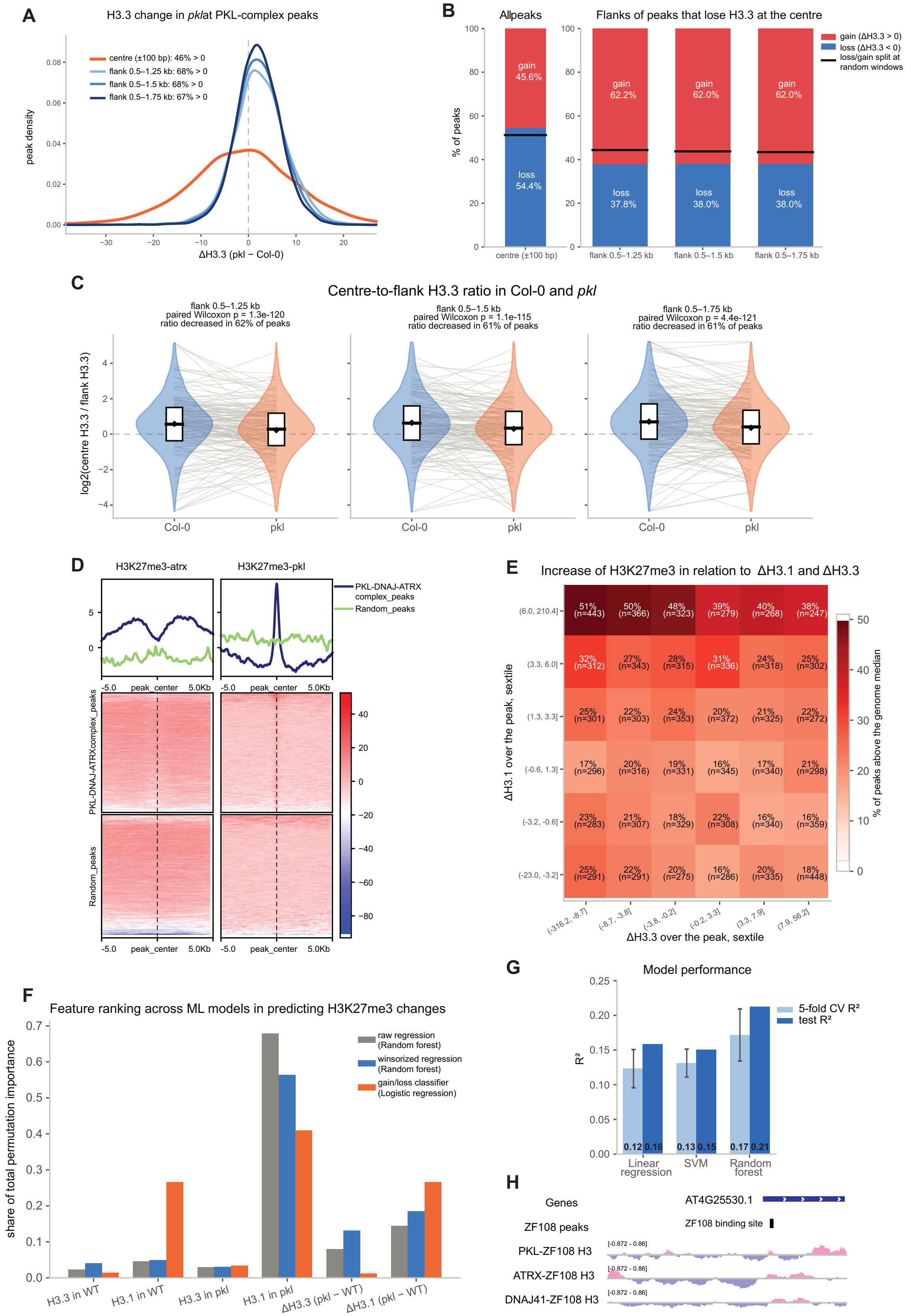
A
H3.3 change in pklat PKL-complex peaks
centre (±100 bp): 46% > 0
flank 0.5–1.25 kb: 68% > 0
flank 0.5–1.5 kb: 68% > 0
flank 0.5–1.75 kb: 67% > 0
peak density
ΔH3.3 (pkl − Col-0)
B
Allpeaks
Flanks of peaks that lose H3.3 at the centre
gain (ΔH3.3 > 0)
loss (ΔH3.3 < 0)
loss/gain split at random windows
% of peaks
gain 45.6%
loss 54.4%
gain 62.2%
loss 37.8%
gain 62.0%
loss 38.0%
gain 62.0%
loss 38.0%
centre (±100 bp)
flank 0.5–1.25 kb
flank 0.5–1.5 kb
flank 0.5–1.75 kb
C
Centre-to-flank H3.3 ratio in Col-0 and pkl
flank 0.5–1.25 kb
paired Wilcoxon p = 1.3e-120
ratio decreased in 62% of peaks
flank 0.5–1.5 kb
paired Wilcoxon p = 1.1e-115
ratio decreased in 61% of peaks
flank 0.5–1.75 kb
paired Wilcoxon p = 4.4e-121
ratio decreased in 61% of peaks
log2(centre H3.3 / flank H3.3)
Col-0
pkl
D
H3K27me3-atrx
H3K27me3-pkl
PKL-DNAJ-ATRX complex_peaks
Random_peaks
-5.0
peak_center
5.0Kb
PKL-DNAJ-ATRXcomplex_peaks
Random_peaks
E
Increase of H3K27me3 in relation to ΔH3.1 and ΔH3.3
ΔH3.1 over the peak, sextile
ΔH3.3 over the peak, sextile
% of peaks above the genome median
(6.0, 210.4]
(3.3, 6.0]
(1.3, 3.3]
(-0.6, 1.3]
(-3.2, -0.6]
(-23.0, -3.2]
(-316.2, -8.7]
(-8.7, -3.8]
(-3.8, -0.2]
(-0.2, 3.3]
(3.3, 7.9]
(7.9, 56.2]
51% (n=443) 50% (n=366) 48% (n=323) 39% (n=279) 40% (n=268) 38% (n=247)
32% (n=312) 27% (n=343) 28% (n=315) 31% (n=336) 24% (n=318) 25% (n=302)
25% (n=301) 22% (n=303) 24% (n=353) 20% (n=372) 21% (n=325) 22% (n=272)
17% (n=296) 20% (n=316) 19% (n=331) 16% (n=345) 17% (n=340) 21% (n=298)
23% (n=283) 21% (n=307) 18% (n=329) 22% (n=308) 16% (n=340) 16% (n=359)
25% (n=291) 22% (n=291) 20% (n=275) 16% (n=286) 20% (n=335) 18% (n=448)
F
Feature ranking across ML models in predicting H3K27me3 changes
share of total permutation importance
raw regression (Random forest)
winsorized regression (Random forest)
gain/loss classifier (Logistic regression)
H3.3 in WT
H3.1 in WT
H3.3 in pkl
H3.1 in pkl
ΔH3.3 (pkl − WT)
ΔH3.1 (pkl − WT)
G
Model performance
R²
5-fold CV R²
test R²
0.12 0.16
0.13 0.15
0.17 0.21
Linear regression
SVM
Random forest
H
Genes
AT4G25530.1
ZF108 peaks
ZF108 binding site
PKL-ZF108 H3
ATRX-ZF108 H3
DNAJ41-ZF108 H3
[-0.872 - 0.86]

**Figure S5. H3 variants and H3K27me3 change concordantly in the *pkl* mutant. A.** Density plot showing the distribution of H3.3 changes (*pkl* minus Col0) at the center and flanking regions of PKL-DNAJ-ATRX complex peaks. **B.** The left stacked bar plot illustrates the percentage of PKL-DNAJ-ATRX complex peaks that gain or lose H3.3 at the peak center. The four bars on the right show, among peaks with decreased H3.3 at the center, the percentage of these peaks that display either H3.3 gain or loss at the flanking regions. **C.** Paired violin plots showing the H3.3 center-to-peak ratio in Col0 and *pkl* mutant across three definitions of flanking regions (0.5-1.25kb, 0.5-1.5kb, and 0.5-1.75kb). **D.** H3K27me3 ChIP-seq signal of *atrx* and *pkl* normalized to Col-0 at PKL peaks and random peaks. **E.** 2D heatmap displaying the increase of H3K27me3 relative to H3.1 and H3.3 changes in the *pkl* mutant. Peaks of the PKL-DNAJ-ATRX complex were categorized based on the changes in H3.1 and H3.3. For each group, the proportion with H3K27me3 changes above the genome-wide median is indicated and color-coded. **F.** Feature importance ranking across 3 ML models. **G.** Random forest vs other regressors performance by 5-fold CV $R^2$ (light, ± SD) and held-out test $R^2$. **H.** Genome browser example at *FWA* loci showing H3 gain introduced by ZF108-tagged PKL, ATRX, and DNAJ41. All ChIP-seq tracks were normalized to Col-0 wild type.

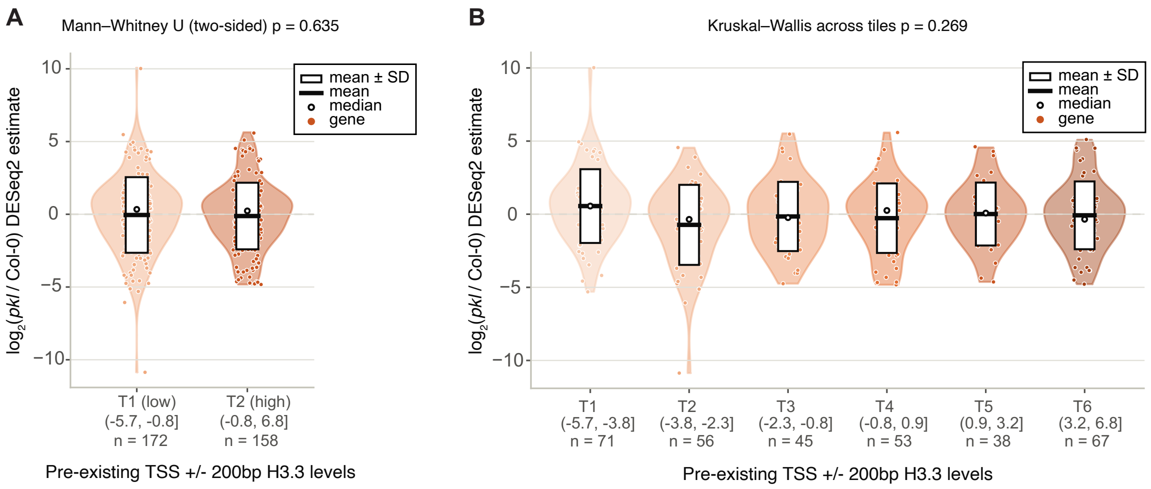


**Figure S6 Expression changes of *pkl* DEGs scale with pre-existing H3.3 only when H3.3 is lost.** RNA-seq log2 fold change (*pkl*/wild-type) of *pkl* DEGs without H3.3 loss at TSSs. *pkl* DEGs are binned into **A**. 2-tiles or **B**. 6-tiles based on the pre-existing H3.3 levels at the TSS of the gene in Col-0, with a cutoff of p-value <0.05. Box indicates mean +/- SD, median is indicated by black dot, and blue points indicate individual genes

**Table S1-12: Summary of AF3 predicted interaction interfaces.** Interaction residue pairs and interface residues of DNAJ41 and PKL (Table S1-2), DNAJ41 and AT1G48560 (Table S3-4), DNAJ41 and AT1G15720 (Table S5-6), DNAJ41 and DF1 (Table S7-8), DNAJ41 and FIP2 (Table S9-10), DNAJ41 and AT2G44730 (Table S11-12).
